# Endophytes induce systemic spatial reprogramming of metabolism in poplar roots under drought

**DOI:** 10.1101/2025.06.02.657501

**Authors:** Jayde Aufrecht, Dusan Velickovic, Robert Tournay, Sneha Couvillion, Vimal Balasubramanian, Tanya Winkler, Daisy Herrera, Robert Stanley, Sharon L. Doty, Amir H. Ahkami

## Abstract

Beneficial endophytes help plants thrive in challenging environments by altering their host’s metabolism, but how these cellular scale metabolic changes propagate to the systems biology scale is unknown. In this work, we employed a high-resolution chemical imaging approach to map metabolic changes at the root zone and cell type levels and found that a 9-strain consortium of beneficial endophytes differentially altered the metabolome of droughted root tissues according to cell types and locations along the root system architecture. Using machine learning (ML) models, we identified root metabolites and exudates that have predictive power over treatment class and could therefore be used as systems biology indicators of drought and endophyte inoculation status. We calculated the correlation between each endophyte and metabolite and found that this relationship shifts under drought conditions, indicating the dynamic role endophytes play in a plant’s microbiome and metabolism in response to environmental changes.

## Introduction

Beneficial microorganisms that reside in the rhizosphere and plant endosphere profoundly influence their host’s health and ability to thrive in suboptimal environments including droughted conditions (1, 2). While the molecular mechanisms underpinning this intricate relationship are well studied in a few key crops, such as rhizobia and soybean, the molecular exchanges and signaling pathways in most other plant-microbe interactions remain unknown. This knowledge gap hinders the translation of laboratory-scale successes in microbial inoculation to agricultural and bioeconomy applications. (3). One school of thought suggests that molecular signals produced by the beneficial microorganisms could prime the plant’s immune response and prepare it for stressful conditions like drought, biotic stresses, or nutrient limitations (4, 5). However, little is known about how microbial induced changes to the plant’s metabolism at the cellular scale could propagate to the whole organism systems biology scale.

Plant root systems are comprised of distinct zones and cell types that define the cell’s role in biological processes. Root-zone specific responses to abiotic stress (6) and exudation profile (7) have been reported. We too recently showed a root zone specific response of allocated C in poplar (8). Within the root zones, the principal specialized functions of each root cell type, *i.e.,* epidermal, cortex, endodermis, xylem, and phloem cells, are described (6, 7, 9, 10). Recent single cell biology studies have suggested novel roles for specific root cell types through identifying a root gene expression atlas in Arabidopsis (11, 12) and tomato (9). While the importance of location-to-function paradigm and the power of this model in revealing new findings in plant responses to environmental perturbations have been reported (13, 14), our understanding of root cell type and zone-specific metabolic changes during plant-microbe interactions (15) and under drought conditions is limited.

Despite numerous studies which report a changing plant metabolome under stress, these metabolomic data have so far had limited application in predicting plant status or physiology (16–20). With the burgeoning use of machine learning (ML) and artificial intelligence (AI) in interpreting and predicting biological functions, it is tacitly assumed that these data science tools will benefit the field of plant biology and microbial interactions as well. Thus far, however, ML and AI models trained on plant molecular data are scarcely reported, likely due to the high cost of molecular data acquisition which precludes sufficiently large datasets needed to train the models (21). Large datasets (i.e. hundreds of samples) may not be strictly necessary if there is a strong relationship between metabolic features and measured plant traits. But whole plant metabolomics risks muddying the relationships between metabolic and plant physiological features since different cell types have been shown to have variable metabolite abundances under changing environment (22).

In this work, we report the impact of inoculation with a 9-strain consortium of beneficial endophyte strains on *Populus trichocarpa*’s intracellular and extracellular metabolic network during water limited conditions but prior to endophyte induced physiological recovery in the plant. With this timeframe, we targeted the plant’s initial responses to drought with and without the endophytes to identify which metabolic pathways and root cell types are most affected by the onset of water stress and how plant-microbe signaling could shift plant metabolism in response. The endophyte strains chosen for this study were isolated from wild poplar and willow trees that were thriving in challenging conditions, and have previously demonstrated drought tolerance as well as N-fixing ability in the host plant *Populus*, making this an ideal model plant-microbe relationship to test (1). Our hypothesis was that this set of endophyte strains would induce host signals under water limited conditions that trigger spatial root metabolic reprogramming and consequentially differential root exudation.

We show that drought and endophyte inoculation have a significant impact on overall root metabolism that is cell type and zone specific. Using cell-type specific metabolomes to train ML models, we successfully predicted plant status (i.e. inoculation and/or drought) based on metabolomic data alone. The metabolites selected by the ML model, as strong indicators of plant status, provide hints at the molecular mechanisms in play.

## Results

### The internal root metabolome shifts according to root zones and cell types in response to drought and endophyte inoculation

Plant morphology and photosynthesis parameters including CO_2_ assimilation and stomatal conductivity clearly showed the effect of drought in stressed plants compared to well-watered controls (Supplementary Figure 1). Endophyte treatment, which in published work has shown benefit to the plant under water limitation using an overlapping subset of endophyte community (1), had not yet recovered the droughted phenotype at the time of harvest in this study (see Supplementary Figure 1). With this timeframe, and with a smaller consortium of strains chosen from that work as well as more diazotrophic strains and synergistic partners (23), we targeted the initial stages of metabolic reprogramming that were likely induced by endophytes during drought onset.

Using Matrix-Assisted Laser Desorption Ionization Mass Spectrometry Imaging (MALDI-MSI), we mapped the relative abundance of 722 putative metabolites across different root zones and cell types from the four treatment conditions: 1. Well-watered control (WWC), 2. Well-watered and endophyte inoculated (WWE), 3. Droughted control (DC), or 4. Droughted and endophyte inoculated (DE). While no metabolites were unique to only one of the four treatments, there were 218 metabolites which were significantly different (one-way ANOVA test, p <0.05) in abundance across the four treatments (Figure. 1A) (see Supplementary Data 1 for all significant metabolites and their p values). For spatial root metabolomic analysis, the longest root originating from the stem base was harvested and subdivided into three main growing regions: the root tip (2cms from the root tip representative of root meristematic zone), root middle (immediate 5 cm above the tip region representative of root elongating zone) and root top (immediate 5 cm above the mid region representative of root differentiating zone). For MALDI-MSI, cross sections were made for root top and middle zones, while longitudinal sections were made for root tips. Root tissue sampled from the top of the root system architecture showed the most significant metabolic changes across treatments within three distinct root cell types (i.e. epidermis, cortex, and vascular) compared to root tissue sectioned from the middle root or the root tip (Figure 1). Drought and endophyte treatments both altered the plant metabolome compared to a well-watered control with 1.0 - 15.5% of the measured metabolome shifting depending on the root cell type and root zone.

**Figure 1.**
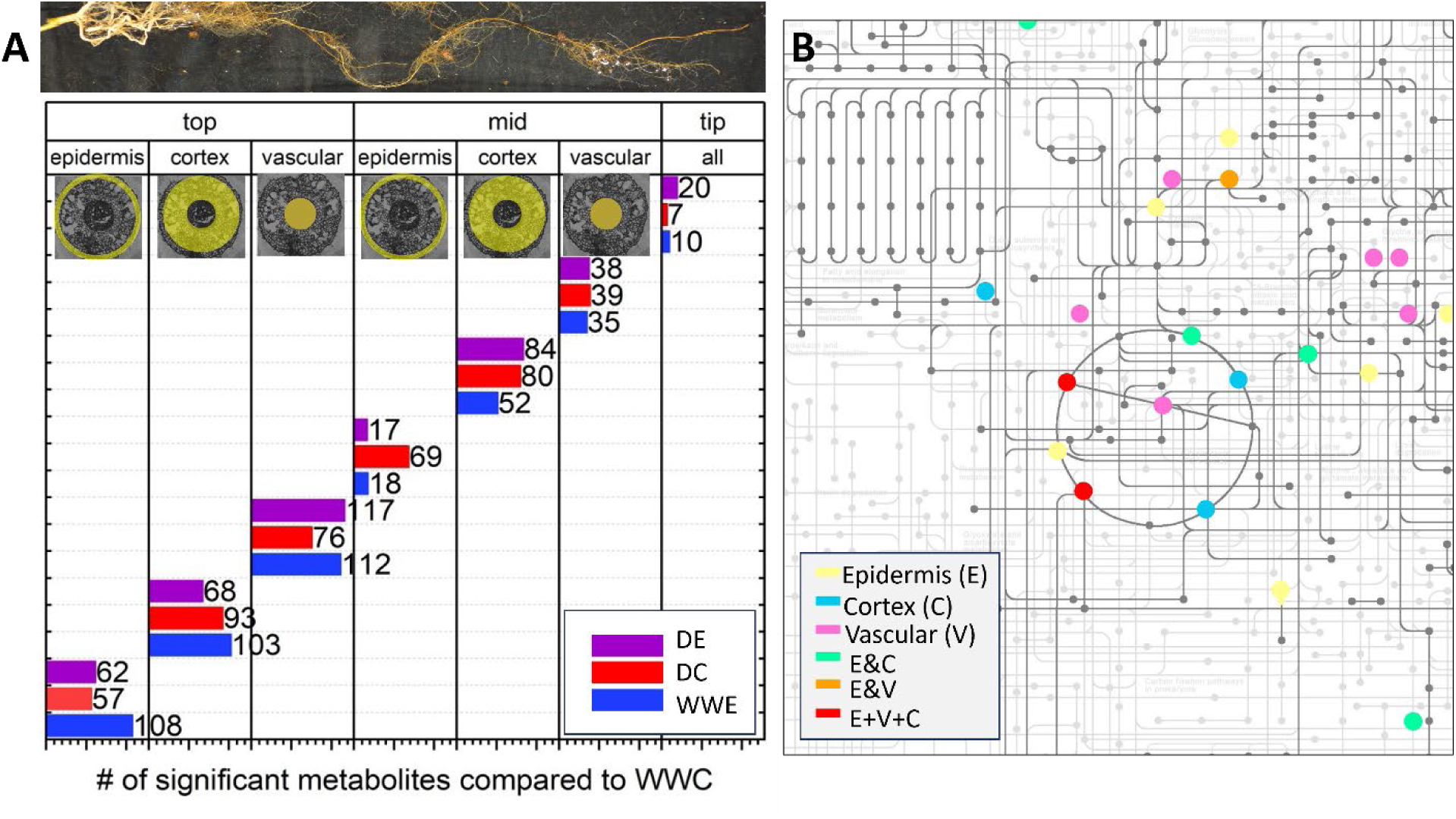
**A)** The magnitude of metabolic shift within root tissue cross sections under drought + endophyte (DE), drought + control (DC), or well-watered + endophyte (WWE) treatments compared to a well-watered control (WWC) depending on the location of the root section (top, mid, tip) and the cell type (epidermis, cortex, vascular) measured. **B)** Endophyte treatment significantly altered several metabolic pathways including the metabolites that belong to TCA cycle as highlighted in the metabolic map of *Populus trichocarpa.* Colored dots indicate metabolites which are differentially abundant in all endophyte treatments (WWE and DE combined) compared to all non-inoculated treatments (WWC and DC combined). Each dot represents one metabolite, and colors indicate in which cell types those metabolites were significantly different as indicated by the legend.

The putative metabolites differentially produced during endophyte treatments (WWE and DE combined) compared to non-inoculated controls (WWC and DC combined) were mapped onto the metabolic network for *Populus trichocarpa.* The most differentially abundant metabolites in each cell type under endophyte treatment were: 12-hydroxydodecanoic acid in the vascular tissues (168.24-fold), 12-hydroxydodecanoic acid in the cortex (253.29-fold), and Sinensetin in the epidermal tissues (257.79-fold). This analysis also revealed that several of the differentially abundant metabolites were involved in the citric acid cycle (Figure 1B, see Supplementary Fig. 2 for full metabolic map). However, the impact of endophyte treatment (i.e. both DE and WWE together compared to WWC and DC together) on the root metabolome was not uniform across cell types; epidermal, cortical, and vascular cells showed different patterns of metabolic shifts under endophyte treatments. For example (iso citrate was reduced in epidermal (0.52-fold) and cortical cells (0.57-fold) in all inoculated treatments; cis-aconitate (0.65-fold) and oxoglutarate (0.46-fold) were decreased only in cortical cells and fumarate was only altered in the epidermis (0.62-fold). These results show the impact that endophytes can have on root metabolism irrespective of environmental condition and act as a baseline when considering the impact of endophytes on droughted plants only.

### Endophyte treatments reprogram root metabolic pathways under drought

We identified 166 features (i.e. metabolites in specific root zones or cell types) that were significantly different when comparing DE vs. DC plants (See supplementary Data 5 for full list). These 166 features matched to 89 potential pathways in the *Populus trichocarpa* metabolome which could be significantly affected by the presence of endophytes (see Supplementary Figure 3 for metabolic map). Of these 89 pathways, 24 had more than one differentially abundant metabolite and were therefore more likely to be affected due to the presence of endophytes (Supplementary Table S1) under drought. Primary metabolic pathways such as the citric acid cycle, reductive citrate cycle, and photorespiration, potentially had increased activity under endophyte inoculation due to the higher increased abundance of related pathway metabolites (compared to other pathways). Some secondary metabolic processes responsible for biosynthesis and degradation of phenolic compounds and fatty acids such as cumate degradation, cutin, suberin and wax biosynthesis, and the shikimate pathway also had potential increased activity due to an increase in associated metabolites in inoculated plants under drought (DE vs. DC treatments) (Supplementary Table S1).

### Endophytes affect the spatial distribution of primary metabolites within the citric acid cycle under drought condition

As shown in Supplementary Table S1, the citric acid cycle was among the metabolic pathways that was significantly altered by the presence of the endophytes under the drought condition. To better understand the magnitude that endophyte inoculation has on the central metabolism (specifically the citric acid cycle) of root cells under drought, we compared representative molecular images of mid root cross sections from two droughted plants: one inoculated with endophytes (DE) and one control (DC) (Figure 2). The non-inoculated roots showed increased abundance in metabolites in all cell types through the first three steps of the citric acid cycle compared to the DE roots (average DE/DC fold changes, (iso)citrate = 0.79, cis-aconitate =0.72). After iso-citrate and before the split to the glyoxylate pathway, the metabolites shifted to higher abundance in the DE root epidermis compared to the DC root epidermis (average DE/DC fold changes, succinate = 1.1, fumarate =1.5, malate = 1.92)(Supplementary Data 2).

**Figure 2.**
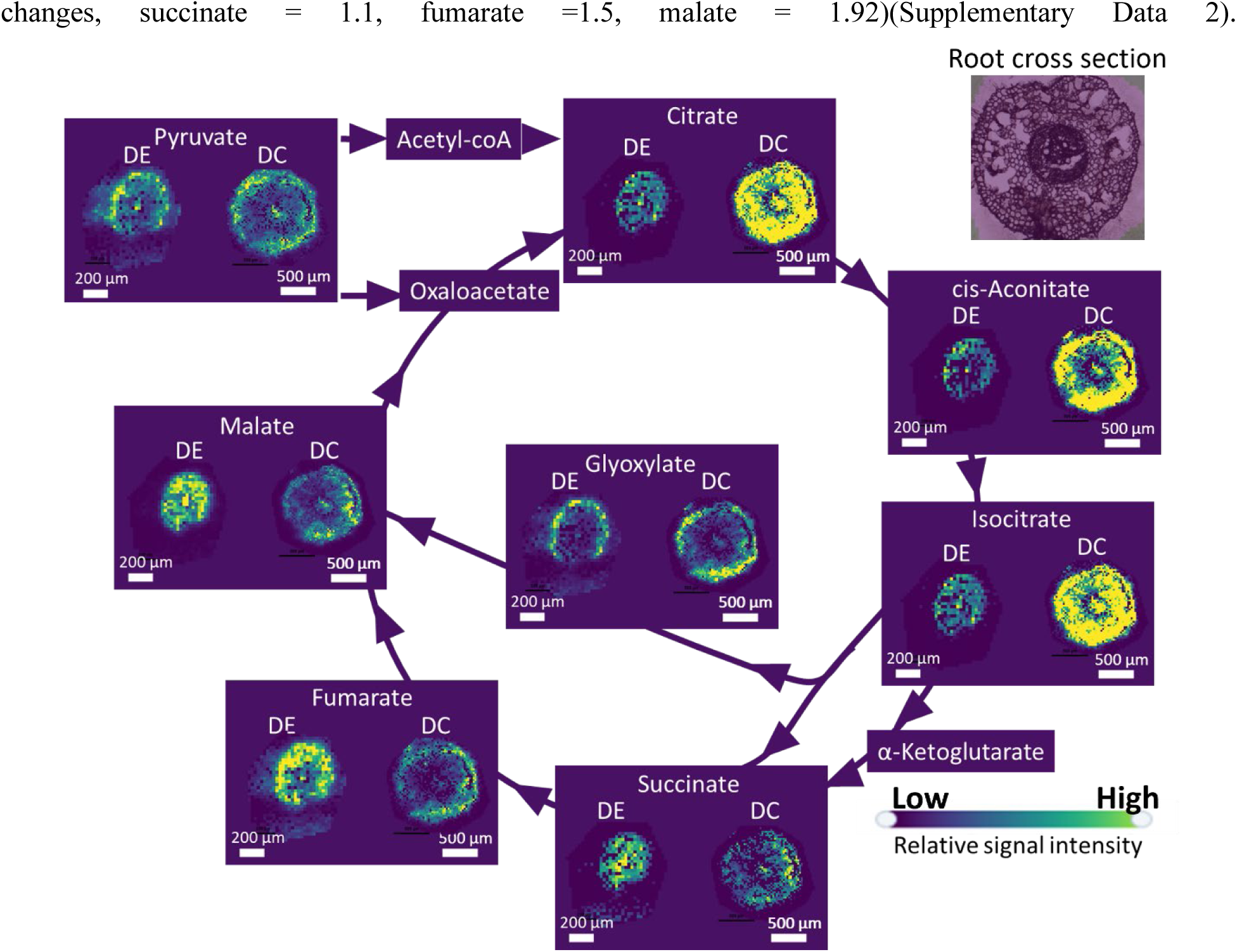
Representative molecular images of root cross sections from two plants (one drought + endophyte (DE) treated plant and one drought + control (DC) treated plant) show that the abundance and spatial distribution of each metabolite in the citric acid cycle is affected by endophyte treatment. Note that relative abundances may only be compared between images of the same metabolite, and the metabolic maps of isocitrate and citrate are the same since the MALDI-MSI cannot distinguish between these isomers.

### Root exudate profiles shift under drought and endophyte treatments

As drought and endophyte inoculation were found to alter the internal root metabolome, we next examined whether metabolites exuded by the roots were also altered under these same treatments. Using liquid-chromatography mass spectrometry (LCMS), we identified 338 metabolites in root exudates from plants across the four treatments (Figure 3A). The metabolites which shifted the most in DE plants compared to the WWC plants were mostly secondary metabolites which ranged from 64% to 145% abundance compared to the average value (normalized to 100% here) of the WWC treatment group (see Supplementary Data 3 for full list of exudates). We compared the root internal spatial metabolome and exuded metabolome and found that only 103 putative metabolites were located both within the roots and within the exudates (see Supplementary Data 4 for full list). However, because MALDI cannot differentiate isomers with the same chemical formula, there may be more metabolites which were present in both the internal and extracellular root metabolome. For these 103 putative metabolites, we compared the average abundance value of the metabolite inside all root tissues (all cell types and root zones) compared to the average abundance value in the root exudate for all treatments and did not see any correlations emerge, even when metabolites where binned by their number of C atoms (Supplementary Figure 4). This finding suggests that the mechanism for metabolite exudation is not exclusive to molecular size and is likely not passively driven by concentration gradients across all root tissues.

**Figure 3.**
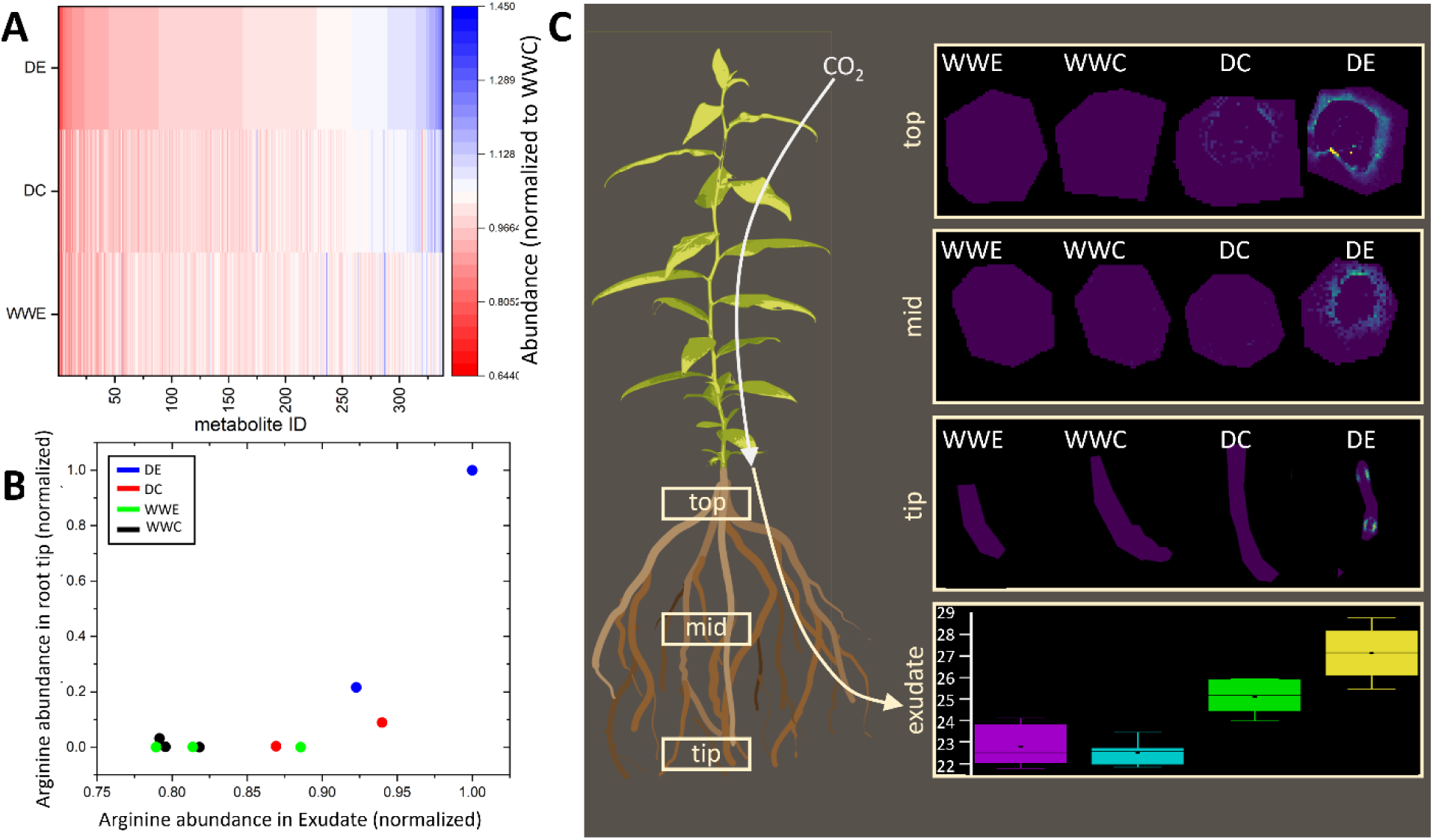
**A)** The root exometabolome was significantly impacted by drought (DC), endophyte (WWE) inoculation, and the combination thereof (DE), compared to a well-watered control (WWC) as shown by the normalized relative abundance of each metabolite. For a full list of metabolites corresponding to the metabolite IDs see Supplementary Data 3 . **B)** Standardized arginine abundances in root tips of all plants showed a significant correlation (Pearson’s coefficient =0.75) to the standardized arginine abundances in the root exudates. Each dot corresponds to a plant replicate. **C)**. Arginine was significantly increased in root exudate (relative abundance shown) and root tip in DE conditions. Arginine was also significantly accumulated in root middle and top zones, specifically in epidermis cells, under DE treatment.

One primary metabolite, the amino acid arginine, was significantly increased (45% increase) in root exudates under the DE treatment compared to the WWC condition. This finding led us to examine the distribution of arginine within the internal root metabolome. We found that arginine levels were significantly higher in root tip longitudinal sections and epidermal cells in top and middle root zones in DE plants compared to all other treatments (Figure 3C). Although, like all other metabolites, arginine also did not show a correlation between its abundance in all root tissues and abundance in the root exudate, we did see a correlation between the abundance of arginine in the root tips specifically and the abundance in the root exudate (Figure 3B, Pearson’s coefficient = 0.75). In fact, arginine was the only metabolite which demonstrated such a correlation (Pearson’s coefficient > 0.5) between root tip abundance and root exudate abundance across all plants and treatments. This result is in direct contrast with some current theories which posit that metabolites are primarily exuded by root tips (7).

### Machine learning models use indicator metabolites to predict plant treatment status

Given that the endophyte and drought treatments had induced complex, non-uniform changes in the internal and external metabolome of *Populus trichocarpa*, we attempted to determine whether we could use the metabolome information alone to predict whether a given plant had been inoculated with endophytes or whether it had been subjected to drought conditions. We separated each metabolite abundance by its location to the root (top, mid, tip, exudate) and cell type (epidermis, vascular, cortex, exudate) which resulted in 6900 metabolite features (exudates were considered as external location to root). We used lasso regularization to reduce the number of features to 7-10 (see Table 2 and Table 3) which also eliminated any multi-collinearity effects between features (24).

Next, we split the data into train (67%) and test (33%) sets to train and test two machine learning classification models where the classifier labels were (1) either inoculated or not-inoculated and (2) either droughted or well-watered. For each model, we tested three ML approaches: logistic regression, KNN classification, and decision trees. KNN models performed the best with 100% accuracy in ability to predict whether a plant was droughted or whether it had been inoculated with beneficial endophytes (Figure 4). To confirm that we were not overfitting the models, we also used a bagging ensemble method to bootstrap the training datasets and still achieved 100% prediction accuracy.

**Figure 4.**
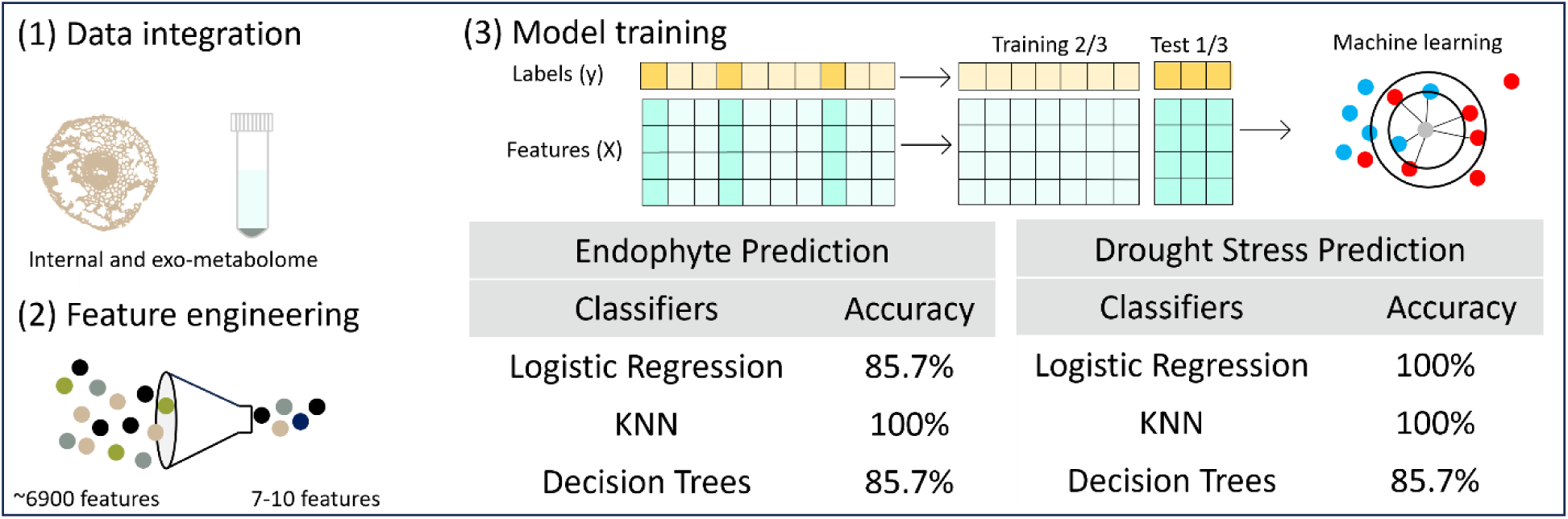
2 ML models were employed to predict the status of each plant (droughted vs. well-watered or inoculated vs. control) based on the metabolome (including internal root tissue and exudates) alone. Metabolite features were reduced from 6900 to 7-10 using lasso regularization feature engineering then split into train and test datasets for three ML approaches.

For both ML models (drought prediction and endophyte prediction), the features selected were a combination of internal and exuded metabolites, with the majority being secondary metabolites (except for arginine, phenylalanine, glutaric acid and succinate). Because these metabolites have demonstrative predictive power over determining the plant’s status, we can consider them to be “indicator molecules” of each condition (i.e. drought or endophyte inoculated). The specific role that each indicator molecule has in regulating plant metabolism under a changing environment is not always fully determined due to a lack of published research on these compounds. However, for each indicator molecule, we have assembled a list of proposed mechanisms by which the molecular compound could be regulating the plant system under each treatment (Tables 2 and 3). These ML-based identified “indicator molecules” could be used in future work to predict plant status and to further unravel molecular mechanisms underlying plant-microbe interactions under water limited conditions.

### The correlation between endophyte and metabolite abundances shifts with drought treatment

To validate endophyte colonization, we quantified bacterial abundance in *P. trichocarpa* roots via ddPCR using previously validated strain-specific primers targeting eight bacterial endophytes: *Herbiconiux* sp. 11R-BC, *Azospirillum* sp. 11R-A, *Sphingobium* sp. 11R-BB, *Sphingobium* sp. WW5, *Rahnella aceris* WP5, *Azotobacter beijerinckii* SherDot1, *Rhizobium* sp. PTD1, and *Rahnella aceris* R10. Analysis excluded the yeast strain *Rhodotorula graminis* WP1 due to primer specificity constraints. Root tissue from non-inoculated plants was pooled together (22 plants) and none of the target endophytes were detected in this sample. Inoculated plants (17 plants) were tested individually for colonization. All eight strains were detected in root tissues of inoculated plants to varying densities (**Table 3**). Detection frequencies varied: PTD1 and 11R-BB colonized all 17 plants; WP5 colonized 16; 11R-BC, WW5, and R10 each colonized 15; 11R-A colonized 14; and SherDot1 colonized 7. Mean colonization densities ranged from 108.4k ±SD 168.2k copies g⁻¹ (11R-A) to 331.8k ± 721.3k copies g⁻¹ (WW5), spanning five orders of magnitude (0.463k–2,996k copies g⁻¹).

Colonization efficiency did not necessarily lead to greater colonization density. WW5 and WP5 demonstrated highest mean densities (331.8k ±SD 721.3k and 268.2k ±SD 547.3k copies g⁻¹, respectively) despite colonizing less than all of the roots, whereas PTD1 and 11R-BB were detected in all roots and had intermediate densities (182.3k ±SD 326.1k and 162.7k ±SD 190.7k copies g⁻¹, respectively). SherDot1 displayed lowest detection frequency (41%) yet had a mean colonization density (166.1k ±SD 133.4k copies g⁻¹) exceeding that of more frequently detected strains. Mean strain detection per droughted root was 7.3 ± 0.85, which was one strain more than the well-watered plant average (6.3±0.81). Colonization densities also had substantial inter-root variation (mean: 1,364k ±SD 2,285k copies g⁻¹), with similar colonization patterns across strains within individual roots. Roots exhibiting low total bacterial loads (e.g. plant IDs: 30, 42, 43) displayed consistently reduced densities across multiple strains, whereas highly colonized roots (e.g. plant IDs: 27, 40, 41) supported elevated densities for most detected strains. This coordinated pattern suggests that root-specific factors may be influencing overall colonization capacity rather than strain-specific competitive interactions. The strain vs. strain correlation matrices corroborate this theory since no negative correlations between strains were observed in droughted plants and only one strain (SherDot1) had a negative correlation with other strains in well-watered plants (**Figure 5**). Overall, these data confirm successful establishment of all consortium members within drought-stressed *P. trichocarpa* roots. Observed colonization densities (0.463k–2,996k copies g⁻¹) align with ranges reported for endophytic bacteria in woody plant tissues(45, 46), though the substantial variation warrants further investigation into host-microbe interaction dynamics under stress conditions.

**Figure 5.**
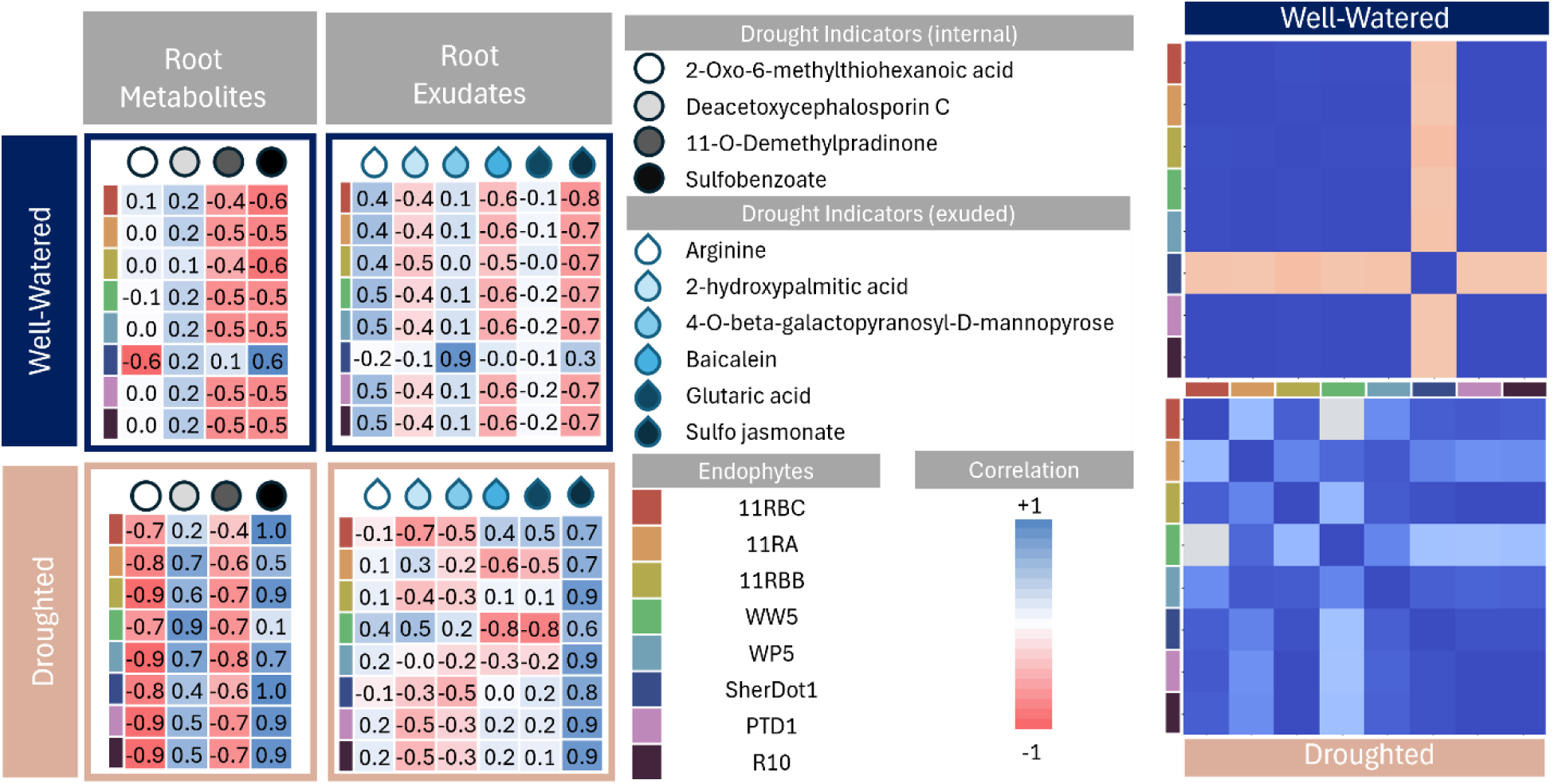
**(Left)** The correlation (Pearson’s coefficient) between each strain abundance and each of 10 drought indicator metabolites was calculated for well-watered and droughted plants. **(Right)** Similarly, the co-correlation matrix between each of 8 endophyte strains living within root tissue is shown for droughted and well-watered plants.

To elucidate the potential role that each endophyte plays during drought stress, we compared the abundance of each strain to the abundance of the drought indicator metabolites identified by our ML model, listed in Table 2. Because the endophytes were measured from bulk root tissue, we averaged the abundance of the drought indicator metabolites within all cell types for mid and top root tissue to create a pseudo-bulk tissue value for each metabolite. The correlation (Pearson’s coefficient) between each strain and each metabolite was calculated for well-watered or droughted plants (Figure 5). For internal root metabolites, the relationships between strain colonization and drought indicator abundances were generally strengthened (either positively or inversely) under drought conditions suggesting that the endophytes have a direct or indirect impact on the metabolites’ abundances either through production and/or consumption or by signalling the plant to alter its metabolism.

The relationships between strain colonization and exudate abundances were generally mixed with some strain-metabolite correlations becoming stronger and others weakening under drought (Figure 5). Notable among the exuded drought indicators are arginine and sulfo-jasmonate. All endophytes (except SherDot1) were inversely correlated to sulfo-jasmonate in well watered plants, suggesting that the endophytes were consuming sulfo-jasmonate or otherwise influencing the plant system to produce less in well-watered conditions. However under drought, the relationship between sulfo-jasmonate and all endophyte strains becomes positively correlated suggesting that the microbes are producing sulfo-jasmonate under drought or otherwise activating the plant hormone biosynthesis pathways to produce more of this hormone. For arginine, however, the relationship between endophytes colonization and metabolite abundance weakens under drought, suggesting that arginine exudation is likely plant-regulated rather than microbial influenced.

## Discussion

It has been proposed that beneficial microbes may prime a plant’s immune system and prepare the host to withstand stressors, but the exact cascade of signals that would induce this systemic change to the plant is unknown. In this work, we targeted the metabolome and exometabolome of roots, as the site of water uptake and endophyte entry, to understand the impact of endophyte inoculation and drought on plant metabolism. We found that the drought stress and endophyte inoculation both significantly and spatially alter a plant’s metabolism. Because no metabolites were unique to the endophyte inoculated plants, we can assume that all identified metabolites are produced, in part, by the plant host, although the endophytes themselves may contribute to the metabolite’s abundance in inoculated plants, and indeed many strains show a strong correlation between colonization and metabolite abundance.

The 9 strains chosen for the consortium were selected to optimize the benefits to plant health under abiotic stresses. Bacterial strains PTD1, WP5, and WW5, and the yeast strain, WP1, were members of larger consortia that conferred drought tolerance to perennial rye grass (47), poplar (1), and conifers (48), increased water use efficiency in rice (2), and increased growth and nitrogen fixation in poplar under nutrient-limited conditions (49). SherDot1 (*Azotobacter beijerinckii*) is a particularly strong nitrogen fixing endophyte (unpublished data). The diazotrophic (nitrogen-fixing) strains, WP5 and 11RA, have elevated nitrogenase activity when mixed with the synergistic strains, WW5 and 11RB (11RB-B and 11R-BC) (23). The same consortium used in this study was shown to increase intrinsic water use efficiency (WUE) in poplar (50).

We observed a considerable impact on the citric acid cycle when we inoculated plants under drought stress. Under stressful conditions, like drought, plants are known to decrease CO_2_ assimilation resulting in a carbon starvation state with reduced glycolysis and acetyl-CoA to fuel energy generation from the citric acid cycle (51, 52). We observed reduced CO_2_ assimilation rates in inoculated and droughted plants (Supplementary Figure 1) consistent with these expectations, and we also observed significantly less glucose within droughted plants compared to well-watered (Supplementary Figure 5). Previously published research has shown that despite drought conditions, plants inoculated with a consortia containing strains which overlap with the consortia used in this study are able to maintain biomass productivity (1). One demonstrated endophyte-induced mechanism that contributes to this enhanced plant productivity under drought is the increase in WUE by the plant due to a reduction in stomata size and increase in stomatal conductivity (50, 53). However, this should lead to higher CO_2_ assimilation in inoculated plants which we did not observe at the time of harvest in this work.

*In lieu* of active C-fixation and assimilation, stressed plants begin to break down biomolecules like proteins and lipids to harvest C for energy production (54). In this work we observed a mixed impact of endophyte inoculation on fatty acid abundances within root tissues (Table S2). For example, 16-hydroxypalmitate had increased abundance under endophyte treatment only in top sections of the root in the vascular cell compartment, while 16-oxopalmitate showed reduced abundance in mid sections of the root within the cortical cells. The particular locations of these fatty acids suggest that under endophyte treatment, there is possible transport of fatty acids from C-sources down through the phloem (explaining the increase in vascular tissues) to be catabolized in cortical cells of newer root tissue for possible various purposes including root growth, C storage, or exudation. Fatty acids have recently been shown to be transported in the phloem which supports this theory (55). ML models also identified 2-hydroxypalmitic acid as an indicator metabolite for drought stressed plants.

The resulting acetyl-CoA from lipid degradation may be fueling the citric acid cycle in DE-treated plants. However, we observed a mix in citric acid cycle metabolite abundances in this condition wherein the first three intermediate metabolites (citrate, aconitate, and iso-citrate) had a reduced abundance in DE plants compared to DC and the final three intermediate metabolites (succinate, fumarate, and malate) had an increased abundance in epidermal tissues under DE compared to DC. In our previous work, we showed that at least one poplar-isolated endophyte (*Burkholderia vietnamiensis* WPB, a nitrogen fixing bacteria) produces citric acid under non-nitrogen fixing conditions likely so that nearby cells that are actively fixing nitrogen can readily consume citric acid for energy (56). In the same work, we also showed that this WPB strain lives inside root epidermal cells of elongating root tissues. If the endophytes are consuming citric acid (and iso-citric acid), this may account for the reduced abundance in the first three TCA metabolic intermediates under DE conditions. Our previous work also demonstrated that the N-fixing endophyte within the consortia produces polyamines and amino acids (56) which are bioavailable forms of N for the plant. Although our MALDI analysis was not optimized for detecting common polyamines like spermine, putrescine, and ornithine, which were below our 150Da molecular weight cutoff, we did detect (N1, N12)-Diacetylspermine to be significantly more abundant in root epidermis of DE plants compared to all other conditions. We also identified 14 total amino acids and amino acid derivatives in DE samples, 12 of which were significant compared to the well-watered control. The enhanced production of amino acids and polyamines by the endophytes could be contributing to the increase in the final three TCA intermediates (succinate, fumarate, and malate) as the carbon skeletons of transaminated amino acids enter the TCA cycle as alpha-ketoglutarate (54). Of course, any increase or decrease in intermediates has an anaplerotic effect on the TCA cycle – it will accelerate or decelerate the rate of energy production, but total energy produced is still limited by the acetyl-coA feedstock. Polyamines themselves are also known to reduce plant stress via anti-oxidation (57), but also act in concert with plant hormones like jasmonic acid which regulate the plants systemic response to stress (58). Our MALDI technique did not pick up any jasmonic acid signal inside of root tissues, however, the LC-MS detected a significant decrease of sulfo-jasmonate in root exudates under DE compared to DC conditions, and endophyte abundance was positively correlated with sulfo-jasmonate abundance under drought.

As previously mentioned, we saw the endophyte treatment enhance amino acid abundances such as arginine. Here, we identified a unique spatial shift of arginine abundances exclusively within DE plants in the epidermal cells of top and middle root zones, as well as in the root tip (Figure 3c). Arginine has the highest C:N ratio among all proteinogenic amino acids, which makes it a primary storage and transport form of organic nitrogen in plants. Apart from its significance in protein synthesis, arginine also acts as a precursor for polyamines and nitric oxide, and therefore it plays a crucial role in plant responses to abiotic stresses (59). Interestingly, arginine was also significantly enriched in the root exudate profiles under DE treatments. As a root exudate, arginine is shown to increase nitrogenase (the enzyme responsible for bacterial N fixation) activity in the soil, thereby increasing nitrogen availability under the harsh conditions of arid environments (60). Our ML analysis also identified arginine as a root exudate indicator to predict drought status (Table 2), highlighting an important role of arginine as a key root metabolite and exometabolite during root-microbial interactions under drought conditions.

Despite the many articles which have characterized plant metabolomes under changing environmental conditions, we were only able to find one research paper in which ML tools were used to predict plant status from metabolomics data (61). One challenge of machine learning – which is even more limiting in deep learning analysis – is the large, expensive sample sets required to train and test the models. Here we used machine learning on a sample size of only 21 plants. We were able to generate an accurate machine learning model without overfitting the data using two key approaches: 1. Feature engineering to reduce the number of features (i.e. metabolites in specific root locations) used to train the model from ∼6900 features to fewer features than the number of samples, and 2. An ensemble bootstrap approach that trained many models on different subsets of the training set and aggregated the prediction accuracy from all models together. Our ML model was successful because there were very strong relationships between the selected features and the classes (i.e. endophyte treatment or drought treatment). Using a machine learning approach allowed us to identify which metabolites have predictive power over treatment class and could therefore be used as indicators of that treatment class.

The metabolites used to predict droughted plants were different than those used to predict endophyte inoculated plants, but both sets of metabolites included a combination of root-exuded and root internal metabolites. Interestingly, the root internal metabolite abundances used in prediction were almost entirely (except for Deacetoxycephalosporin C) sampled from the top region of root tissue (representative of root differentiating zone) closest to the stem base. It is unclear why root tissue from the top region of the root architecture has a stronger predictive power of drought or endophyte treatment class, but this result can be used to inform future root sampling work.

We also observed that most metabolites selected by the feature engineering approach were secondary metabolites with the exception of succinate, phenylalanine, arginine, and glutaric acid. Since most plant metabolomics studies have used GC-MS, which is limited in molecular charge to mass ratio and therefore mostly identifies primary and some smaller secondary metabolites, many of the indicator molecules identified in this work have not been previously published or characterized in plant systems. We employed a relatively new MALDI chemical imaging approach to target carboxylated metabolites which gave us high coverage of the root spatial metabolome and allowed us to identify these indicator molecules (62). Because many of these indicator molecules have not been previously identified, little is known about their influence on plant signaling in drought or endophyte treatments. From the available literature, we identified potential signaling pathways and mechanisms upon which the indicator molecules could act including roles as antioxidants, osmoprotectants, antimicrobials, and quorum sensing molecules (Tables 1 and 2). However, these roles will need to be confirmed through additional experimentation in the future. Approximately, half of the indicator molecules chosen by the feature engineering workflow were root-exuded metabolites, indicating a potential significant role of root exudates on root-microbial interactions under drought conditions. This finding is also promising for future identification of plant status in the field by leveraging established non-destructive and *in situ* exudate sampling approaches like micro-dialysis and cuvette sampling (63). Many of the drought indicator metabolites are positively or inversely correlated to specific endophyte strains and this correlation changes depending on whether the plant is well-watered or droughted. These results shed light on the dynamic role that endophytes play in plant microbiome and metabolism under stress.

**Table 1:** Metabolites selected by lasso-regularization which were used in ML models to predict endophyte inoculation and their proposed mechanism of interaction.

| Prediction of Endophyte Inoculation |  |  |  |
| --- | --- | --- | --- |
| Indicator molecule | Metabolite Location | Molecular class | Proposed mechanism |
| Succinate | Internal (Top Root) | Organic acid | 1. Microbial receptor ligand (25)<br>2. Recruitment of beneficial microbes (26) |
| 10-hydroxydecanoic acid | Root Exudate | Fatty acid | 1. antimicrobial and antifungal activities (27)<br>2. inhibition of histone deacetylases (HDACs) (28) |
| 1-epi valienol 7-phosphate | Internal (Top Root) | Phosphorylated cyclitol | 1. antibiotic synthesis intermediate (29) |
| Phosphoagmatine | Internal (Top Root) | Phosphorylated polyamine | 1. polyamine precursor (30) |
| 3,4, dihydroxybenzoic acid | Root Exudate | Phenolic acid | 1. antimicrobial (31)<br>2. microbial quorum sensing (32) |
| n-octyl sulfate | Root Exudate | Sulfated hydrocarbon | *Unknown* |
| L-Phenylalanine | Root Exudate | Amino acid | 1. shikimate pathway precursor<br>2. antioxidant precursor (33) |

**Table 2:** Metabolites selected by lasso-regularization which were used in ML models to predict drought status and their proposed mechanism of interaction.

| Prediction of Drought |  |  |  |
| --- | --- | --- | --- |
| Indicator molecule | Metabolite type | Molecular Class | Proposed mechanism |
| Arginine | Root Exudate | Amino acid | 1. Polyamine and antioxidant synthesis precursor (34)<br>2. Osmoprotectant precursor (35) |
| 4-sulfobenzoate | Internal (Top Root) | Sulfated organic acid | *Unknown* |
| 2-hydroxypalmitic acid | Root Exudate | Fatty acid | 1. Cutin production and cell turgor |
| 11-O-Demethylpradinone I | Internal (Top Root) | Quinone | *Unknown* |
| 4-O-.beta.-Galactopyranosyl-D-mannopyranose | Root Exudate | Disaccharide | 1. Osmoprotectant (36) |
| Deacetoxycephalosporin C | Internal (Mid Root) | Antibiotic precursor | *Unknown* |
| Baicalein | Exudate | Flavonoid | 1. Antioxidant (37)<br>2. cell turgor (38) |
| 2-Oxo-6-methylthiohexanoic acid | Internal (Top Root) | Carboxylic acid | 1. antioxidant precursor (39, 40) |
| Glutaric acid | Root Exudate | Organic acid | 1. osmoprotectant (41) |
| Sulfo jasmonate | Root Exudate | Hormone derivative | 1. antioxidant enhancement (42)<br>2. promote stress reducing metabolism (43) |
|  |  |  | 3. regulation of stomata and CO<sub>2</sub> assimilation (42) |
|  |  |  | 4. phytohormone crosstalk (44) |

**Table 3.** Colonization densities (copies g-^1^ root tissue) of eight bacterial endophyte strains across 17 root samples, with mean and std. dev. (SD). Strains: *Herbiconiux* sp. l !R-BC, *A:;ospirillum* sp. l JR-A, *Sphingobium* sp. l lR-BB, *Sphingobiwn* sp. WW5, *Rahnella aceris* WP5, *A=otobacter beijerinckii* SDl, *Rhi:obium* sp. PTDl, and *Rahnella aceris* RlO. “nd” indicates target DNA below detection limits. Well-watered control (WWC), Well-watered and endophyte inoculated (WWE), Droughted control (DC), Droughted and endophyte inoculated (DE). Colonization efficiency represents proportion of roots with detectable strain presence. Mean and SD calculated from positive detections only. **All copies g-^1^values XlO00. Colonization efficiency** = **n/17**

| Plant ID | Treatment | 11R-BC | 11R-A | 11R-BB | WW5 | WP5 | Sher Dot1 | PTD1 | R10 |
| --- | --- | --- | --- | --- | --- | --- | --- | --- | --- |
| Pooled 1-5, 7-14, 17-21, 23-26 | WWC and DC | nd | nd | nd | nd | nd | nd | nd | nd |
| 27 | WWE | 419.4 | 660.0 | 751.4 | 2996 | 2303 | nd | 1364 | 1293 |
| 28 | WWE | nd | 43.8 | 156.6 | 166.6 | 29.4 | nd | 27.0 | nd |
| 29 | WWE | 15.2 | 18.7 | 64.5 | 285.3 | 36.0 | nd | 84.6 | 88.6 |
| 30 | WWE | 82.7 | nd | 252.5 | 321.2 | 333.6 | nd | 124.0 | 331.9 |
| 31 | WWE | nd | 5.9 | 4.8 | nd | 1.3 | nd | 0.463 | 4.6 |
| 32 | WWE | 27.3 | 11.2 | 21.8 | 15.8 | 5.8 | nd | 26.8 | 36.7 |
| 33 | WWE | 32.2 | 46.0 | 36.3 | 63.5 | nd | 40.0 | 49.5 | 51.2 |
| 34 | WWE | 83.1 | 79.1 | 71.5 | 78.0 | 80.0 | nd | 95.0 | 95.3 |
| 35 | WWE | 27.6 | 33.3 | 127.2 | 68.1 | 14.3 | nd | 17.6 | nd |
| 36 | DE | 179.1 | 282.0 | 217.5 | 169.8 | 183.4 | 311.1 | 120.2 | 163.6 |
| 37 | DE | 85.2 | 128.1 | 136.2 | 138.0 | 84.1 | 170.0 | 106.1 | 176.8 |
| 38 | DE | 65.8 | 36.0 | 17.2 | nd | 46.2 | nd | 24.7 | 19.4 |
| 39 | DE | 73.7 | nd | 111.6 | 119.8 | 126.7 | nd | 149.9 | 122.6 |
| 40 | DE | 242.0 | 111.1 | 314.9 | 418.4 | 501.3 | 230.8 | 343.3 | 288.9 |
| 41 | DE | 884.3 | 59.5 | 458.0 | 106.5 | 454.7 | 368.5 | 556.8 | 450.9 |
| 42 | DE | 8.4 | 2.5 | 8.5 | 15.5 | 48.4 | 8.1 | 4.1 | 3.2 |
| 43 | DE | 15.0 | nd | 14.5 | 14.0 | 42.8 | 34.0 | 4.3 | 3.6 |
| **Colonization Efficiency** |  | 0.88 | 0.82 | 1.00 | 0.88 | 0.94 | 0.41 | 1.00 | 0.88 |
| **Mean copies g<sup>-1</sup>** |  | 149.4 | 108.4 | 162.7 | 331.8 | 268.2 | 166.1 | 182.3 | 208.7 |
| **±SD** |  | 223.3 | 168.2 | 190.7 | 721.3 | 547.3 | 133.4 | 326.1 | 317.3 |

## Conclusion

In this work, we mapped metabolic changes at the root cell level and found that endophytes spatially alter the metabolome of droughted root cells according to root cell types and zones along the root system architecture. We observed that endophyte activity in droughted plants before the recovery of plant physiology involved a cascade of tens of plant metabolic pathways, providing insights into potential mechanistic interactions for endophyte-conferred drought tolerance. Using a machine learning approach, we identified metabolites that have predictive power over treatment class and could therefore be used as whole systems biology indicators of that class. These indicator molecules could be used in future work to predict plant status in the field and to further target specific plant-microbe interaction mechanisms.

## Methods

### Plant growth, Drought stress and harvesting

*P. trichocarpa* stem cuttings were made from the lateral branches of young healthy plants. Small stem sections containing a nodal leaf at the top of the section were made. The leaf was trimmed down to ∼ 1 square inch and soaked in 1% ZeroTol for a 1-3 minutes. The base of the cuttings were then dipped in Rhizopon AA #2 rooting hormone powder and inserted into pots with Pro-Mix BX without mycorrhizae. The growth medium and cuttings were grown under a humidity dome in humid moist conditions in a growth chamber set to 16h light at 24 C and 8h dark at 20 C and 60% relative humidity. Five weeks later the healthiest rooted cuttings were transplanted into larger pots with the Pro-Mix BX without mycorrhizae. The plants were watered every other day with 100 mL of distilled water. Once a week 100 ppm of Jack’s Professional fertilizer was used instead of water. The first bacterial inoculation occurred 5 weeks after transplanting. Each day after the plants were inoculated the watering was reduced to 50 mL to avoid washing the inoculant from the soil. One week after the first inoculation drought was initiated and the watering volume was reduced 50% for the droughted plants. A second inoculation occurred four weeks after the first inoculation. To induce prolonged drought stress, we first determined the 100% soil water content (SWC) in well-watered pots, from which we calculated a target of 25%-30% SWC. To do this, we harvested a separate set of well-watered pots, dried the soil at 60°C for three days in an oven to measure the dry soil weight, thereby calculating the 100% SWC. We then calculated the required 25%-30% SWC to induce prolonged drought stress. The drought-stressed plants were weighed regularly and watered to maintain this reduced water content. Once we reached the 25%-30% SWC, these conditions were sustained for an additional 5 days to create prolonged water deficit stress. At the end of stress, both well-watered and prolonged drought stressed pots were harvested. The age of plants in the time of harvest was about 15 weeks old. Photosynthesis measurements were done using a Li-COR 6800 in the morning before harvesting the plants as well as phenotype measurements. The roots were gently separated out from the soil by shaking using distilled water and a section was taken for MALDI imaging (see below).

### Exudate collection

The plants were placed in a container with MiliQ water covering the roots. They sat for 2 hours at room temperature for exudate collection. The plant tissue was then harvested, and the root exudate samples were filtered using a at 0.22 μm filter attached to a vacuum pump (64). The exudates were flash frozen in liquid nitrogen and stored at −80 C for processing.

### Endophyte culture and inoculation

Endophyte strains were isolated from wild poplar according to Doty et. al 2009 and closest species were identified through genetic sequencing(65). Poplar plants in this study were inoculated with a synthetic community of 8 isolates: 1. *Rhizobium tropici* (PTD1), 2*. Rahnella aceris* (WP5), 3. *Sphingobium salicis* (WW5), 4. *Rhodotorula graminis* (WP1), 5. *Azotobacter beijerinckii* (SherDot1) 6. *Rahnella aceris* (R10) 7. *Azospirillum sp.* (11R-A), and 8. An isolate referred to as 11R-B that we have since found to consist of two strains: *Sphingobium* sp. (11R-BB) and *Herbiconiux* sp. (11R-BC). Each isolate was individually cultured overnight, centrifuged to pellet cells, and resuspended in phosphate buffer saline to an optical density (OD) at 600nm of 0.0125. Strains were then combined into a total inoculum in which cell concentration was measured to 0.1 OD at 600nm. Endophyte-treatment plants were inoculated at two separate instances to ensure that the endophyte consortium was able to colonize the plant endosphere. The first inoculation instance occurred five weeks after transplanting to pots in which plants were inoculated with either 10mL of inoculum (endophyte group) or phosphate buffered saline (PBS)(control group). The second inoculation instance occurred four weeks after the first inoculation in which plants were inoculated with either 2.5mL of fresh inoculum (endophyte group) or phosphate buffer saline (PBS) (control group).

### Quantification of endophyte colonization using digital-droplet PCR

Poplar roots were ground in liquid nitrogen and stored at –80 °C. Total DNA was isolated from 50–70 ±10% mg tissue using the DNeasy Power Plant Pro Kit (Qiagen, 69206) with modifications: stock UFO beads replaced with six 2.8 mm ceramic beads (Qiagen, 13114-325), 6 min disruption at 30 Hz (tube racks rotated after 3 min), and incubation in prewarmed elution buffer (10 min, 60 °C). DNA purity (A260/280) and concentration were measured using Nanodrop Lite and Qubit 4 fluorometer, respectively. Extracts were stored at –20 °C. Strain-specific primers (SSP) design was described in (50), adapted from (66) and (67). Briefly, endophyte genome assemblies were segmented (300 bp) using BBTools shred.sh. Segmented files were imported into *Geneious Prime* (v2025.0.3) and subjected to sequential BLASTn searches(68) against local genomic databases to systematically eliminate non-unique sequences, retaining segments lacking database matches. Local databases were constructed from concatenated complete genome sequences(69), employing progressive filtering: same-genus databases, closely related genera databases, and *Populus trichocarpa* Nisqually-1 reference genome (GCF000002775.5) to prevent off-target amplification. *Primer3* designed primer sets for unmatched sequences targeting 60-150 bp products(70). Digital PCR products underwent final BLASTn screening against NCBI core nucleotide database, with remaining sequences designated strain-specific primer targets. Reaction mixtures contained 11 μL EvaGreen supermix, 2 μL BSA (0.5 μg/μL), 1 μL EcoRI (10 U/μL), 150 nM each primer, 5 μL template DNA, and nuclease-free water to 22 μL. Droplets generated using QX200 automated droplet generator were amplified: 95 °C (5 min), 40 cycles of 95 °C (30 s) and 52 °C (1 min) with 2 °C/s ramp rate, then 4 °C (5 min), 90 °C (5 min). Positive droplets were quantified using QX200 droplet reader and QX Manager Software v2.1.0. Data acceptance required >10,000 droplets per well and >2-fold fluorescence separation between positive/negative populations. Manual thresholds applied when clear bimodal distribution existed without automatic threshold assignment. Samples failing criteria were classified below detection limits. Controls included no-template controls (NTC), uninoculated plant DNA extracts (NC), and positive controls (PC) using spiked uninoculated DNA with serially diluted isolate genomic DNA (10⁻² to 10⁻⁴). Correlation matrices were calculated using python pandas API.

### MALDI-imaging of root cross-sections

Plants were removed from growing pots, and bulk soil was gently removed from the roots by shaking the roots. The roots were gently washed to remove rhizosphere soil adhering to the roots. The longest root originating from stem base was carefully cut, which was divided into tip region (2cm from root tip), mid region (immediate 5 cm above the tip region) and top regions (immediate 5 cm above mid region). Top, mid and tip pieces of the roots were dissected and immediately embedded in a mixture of 7.5% Hydroxypropyl methylcellulose (HPMC) and 2.5% polyvinylpyrrolidone (PVP)9+, underwent a controlled freeze with liquid nitrogen and kept frozen at −80C in the freezer until cryosectioning. Samples were cryosectioned at −10 °C using a CryoStar NX-70 Cryostat (Thermo Scientific, Runcorn, UK), and 12 µm tissue sections were subsequently thaw-mounted onto indium tin oxide (ITO) coated glass slides. Top and mid root pieces were cryosectioned in the plane to produce cross-sections, while tip root pieces were sectioned in the plane to produce longitudinal sections. All root tissues were imaged within one week of cryosectioning and were stored at −80 °C in vacuum sealed bags with desiccant until analyzed. Root tissues sections were on tissue chemical derivatized (OTCD) using previously developed workflow (62). Briefly, agues solutions of EDC (6mg/mL) and 4-APEBA (2 mg/mL) were sequentially sprayed over the sections using the M-5 sprayer (HTX Technologies, Chapel Hill, NC) at 25 µL/min flow rate, 1,200 mm/min linear flow, at 37.5 °C, at 3 mm track spacing with a crisscross pattern, and 2 s drying period was added between 4 cycles applied, with 10 PSI of nitrogen gas and a 40 mm nozzle height.

All imaging was performed on a Bruker Daltonics 12T solariX FTICR MS, equipped with a ParaCell. This instrument has an Apollo II ESI and MALDI source with a SmartBeam II frequency-tripled (355 nm) Nd: YAG laser (Bremen, Germany). Positive ion mode OTCD acquisitions were acquired with broadband excitation from m/z 98.3 to 1,000, resulting in a detected transient of 0.5593 s— the observed mass resolution was ∼110k at m/z 400. FlexImaging sequences were directly imported into SCiLS Lab (Bruker Daltonics, v.2023.a Premium 3D) using automatic MRMS settings. Ion images were directly processed from the profile datasets within SCiLS Lab, and automated annotation of the centroided dataset was completed within METASPACE with a chemical modifier corresponding to the mass shift expected from 4-APEBA derivatization (+ C18H22N2Br, + 345.09663 Da). KEGG-v1 and BraChemDB-2018-01 were used as metabolite databases for annotations.

### LCMS identification of root exuded metabolites

Metabolites were analyzed using reverse phase (RP) and hydrophilic interaction chromatography (HILIC) separations on a Thermo Fisher Scientific HF-X mass spectrometer (Thermo Scientific, San Jose, CA) coupled with a Waters Acquity UPLC H class liquid chromatography system (Waters Corp., Milford, MA). Metabolites were brought up in 200 uL of 50% LCMS grade methanol and 50% nanopure water. RP separation was performed by injecting 5 uL of sample onto a Thermo Scientific Hypersil GOLD C18 column (3 µm, 2.1 mm ID X 150 mm L) heated to 40°C. Metabolites were separated using an 18 minute gradient with data collected on the first 17 minutes. The RP mobile phase A consisted of 0.1% formic acid in nanopure water and a mobile phase B consisting of 0.1% formic acid in LCMS grade acetonitrile. The gradient used was as follows (min, flowrate in mL/min, %B): 0,0.4,10; 2,0.4,10; 11,0.4,90; 12,0.4,90; 12.5,0.5,90; 13.5,0.5,10; 14,0.5,10; 14.5,0.4,10; 15,0.4,10. Positive and negative ion modes were analyzed in separate injections. HILIC separation was performed by injecting 3 uL of sample onto a Waters Acquity BEH Amide column (130 Å, 1.7 µm, 2.1 mm ID X 100 mm L) heated to 55°C. Metabolites were separated using a 16 minute gradient with data collected for the first 15 minutes. The HILIC mobile phase A consisted of 0.05% ammonium hydroxide, 5% LCMS grade acetonitrile, and 94.95% 10 mM ammonium acetate in nanopure water with a mobile phase B consisting of 0.05% ammonium hydroxide in 99.95% LCMS grade acetonitrile. The gradient used was as follows (min, flowrate in mL/min, %B): 0,0.3,95; 6,0.3,37; 7,0.3,37; 7.1,0.3,95; 7.2,0.5,95; 9.5,0.5,95; 9.7,0.3,95; 16,0.3,95. Positive and negative ion modes were analyzed in separate injections. For both RP and HILIC separations the Thermo Fisher Scientific HF-X was equipped with a HESI source and high flow needle with the following parameters: spray voltage of 3.6 kV in positive mode and 3kV in negative mode, capillary temperature of 300°C, probe heater temp of 370°C, sheath gas at 60 L/min, auxiliary gas at 25 L/min, and spare gas at 2 L/min. Metabolites were analyzed at a resolution of 60 k and a scan range of 50 to 750 m/z for parent ions followed by MS/MS HCD fragmentation which is data dependent on the top 4 ions with a resolution of 15 K and stepped normalized collision energies of 20, 30, and 40.

Metabolite identifications were made using MS-DIAL v4.92 for peak detection, identification and alignment (71). For metabolomics, the experimental data was matched both to in-house libraries (m/z less than 0.003 Da, retention time less than 0.3 min, MS/MS spectral match) and a compilation of publicly available MS/MS databases (m/z less than 0.003 Da, MS/MS spectral match). The tandem mass spectra and corresponding fragment ions, mass measurement error, and aligned chromatographic features were manually examined to remove false positives. Relative quantification was performed by calculating peak areas on extracted ion chromatograms of precursor masses. Statistical analysis of metabolomics data was performed using the PMart web application (72). Data was log2-transformed and normalized via global median centering. Statistical comparisons were performed using ANOVA with a Holm test correction (73).

### Machine Learning Models

The machine learning models were built in Python using the sci-kit learn library (74). The dataframe used to train and test the models was comprised of 21 total samples (i.e. individual plants, 11 well-watered, 10 droughted, 9 inoculated, 12 non-inoculated) and 6900 features (i.e. metabolite abundances in a certain sampled location). First, the metabolite data were split into train and test datasets with 67% of the data allocated to the training dataset and 33% to the test dataset. Next the metabolites abundances were standardized within each feature. A lasso regularization model with an alpha value of 0.1 was fit to the training data and used to reduce the number of features in the final machine learning model (24). Lasso model training error for the endophyte prediction model was 0.018 and Lasso model testing error was 0.137 and resulted in 7 features (see Supplementary Tables 2 and 3 for lasso coefficients for each feature). The lasso model training error for the drought prediction model was 0.015 and the Lasso model testing error was 0.12 and resulted in 10 features. Next the filtered features were isolated from the original dataset to train the machine learning models. The data were again split into 67% training and 33% test sets and standardized. The logistic regression model hyperparameter was 1000 max iterations while the KNN hyperparameter was one nearest neighbor. Once KNN was judged to be the best performing model, an ensemble method was used to test overfitting with 100 estimators in a bagging classifier approach.

## Author contributions

J.A., V.B., A.H. A. conceived and planned the experiments. J.A., V.B., T.W., D.H., R.S. and R.T. carried out the experiments. J.A., D.H., S.C., V.B., R.S., R.T. contributed to sample preparation and analysis. J.A., S.D., A.H. A. contributed to the interpretation of the results. J.A. and A.A. took the lead in writing the manuscript. A.H.A and S.D. acquired funding and provided the endophyte strains. All authors provided critical feedback and helped shape the research, analysis and manuscript.

## Acknowledgements

This work was supported by funding from the Biological and Environmental Research (BER) in the U.S. Department of Energy (DOE) Office of Science, (Endo*Populus* project, Award Number: DE-SC0021137). Part of this work was conducted at EMSL (Environmnetal Molecular Sciences Laboratory), a DOE Office of Science User Facility sponsored by the Office of Biological and Environmental Research and operated under Contract No. DE-AC05-76RL01830, located at Pacific Northwest National Laboratory (PNNL).

## Supplementary Files/Tables/Figures

Supplementary Data 1: List of metabolites which were significantly different in abundance across all 4 treatments and their corresponding ANOVA test p values.

Supplementary_Data_2: MALDI chemical imaging data for root internal metabolomics

Supplementary Data 3: List of exuded metabolites and their abundances normalized by the average abundance in the WWC treatment group.

Supplementary Data 4: List of metabolites which were located both inside the root (detected by MALDI-MSI) and in the exudate fraction (detected by LCMS). Integration of the two mass-spectrometry datasets was done by comparing the chemical formula of each metabolite and using the LCMS data to identify metabolite name.

Supplementary Data 5: List of 166 features (i.e. metabolites in specific root locations or cell type) that were significantly different between DE and DC plants.

Supplementary Data 6: Raw digital-droplet PCR data showing samples, targets, concentration, status, and accepted, positive, and negative droplets.

Supplementary Data 7: Concentration of strain-specific bacterial endophytes and estimated colonization density in *Populus trichocarpa* root samples. For each sample, the table presents total root mass, DNA extraction and quality metrics, and digital droplet PCR quantification of eight endophyte strains: Herbiconiux sp. 11R-BC, Azospirillum sp. 11R-A, Sphingobium sp. 11R-BB, Sphingobium sp. WW5, Rahnella aceris WP5, Azotobacter beijerinkii SD1, Rhizobium sp. PTD1, and Rahnella aceris R10. Concentrations (copies/uL reaction) and estimated colonization densities (cells/g root tissue) were determined by strain-specific primers and digital droplet PCR. “NA” indicates no amplification detected for the corresponding strain in the sample.

**Supplementary Fig. 1.**
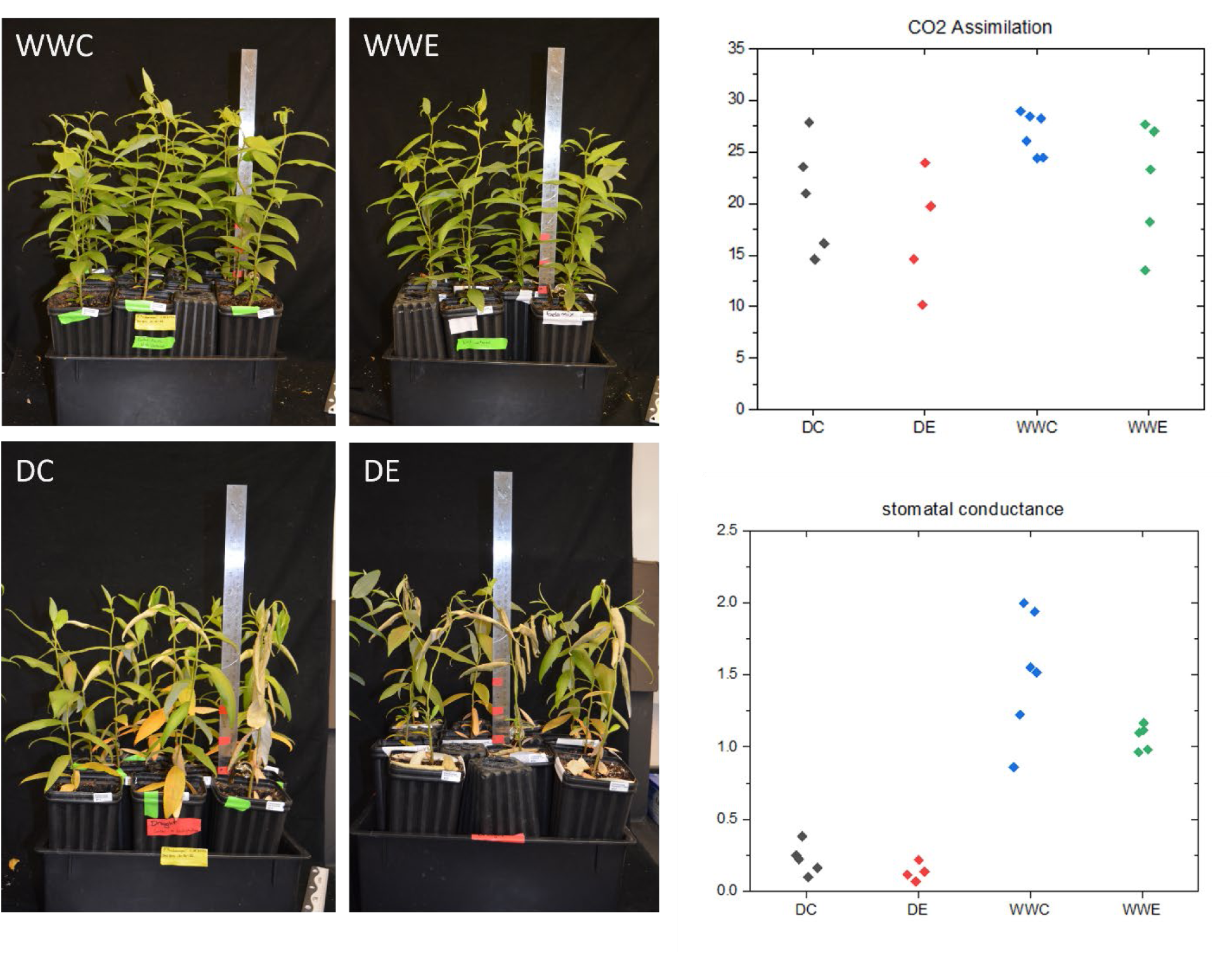
*P.trichocarpa* plants were inoculated with endophytes and exposed to drought stress conditions before used for phenotyping and physiological measurements. (**Left**) Photographs within each treatment group: droughted control (DC), droughted and endophyte inoculated (DE), well-watered control (WWC), and well-watered and endophyte inoculated (WWE) showing poplar plant morphology in control and treated conditions on the day of harvest. (**Right**) Measurements of stomatal conductance and CO2 assimilation was performed on growing plants pre-harvest.

**Supplementary Figure 2:**
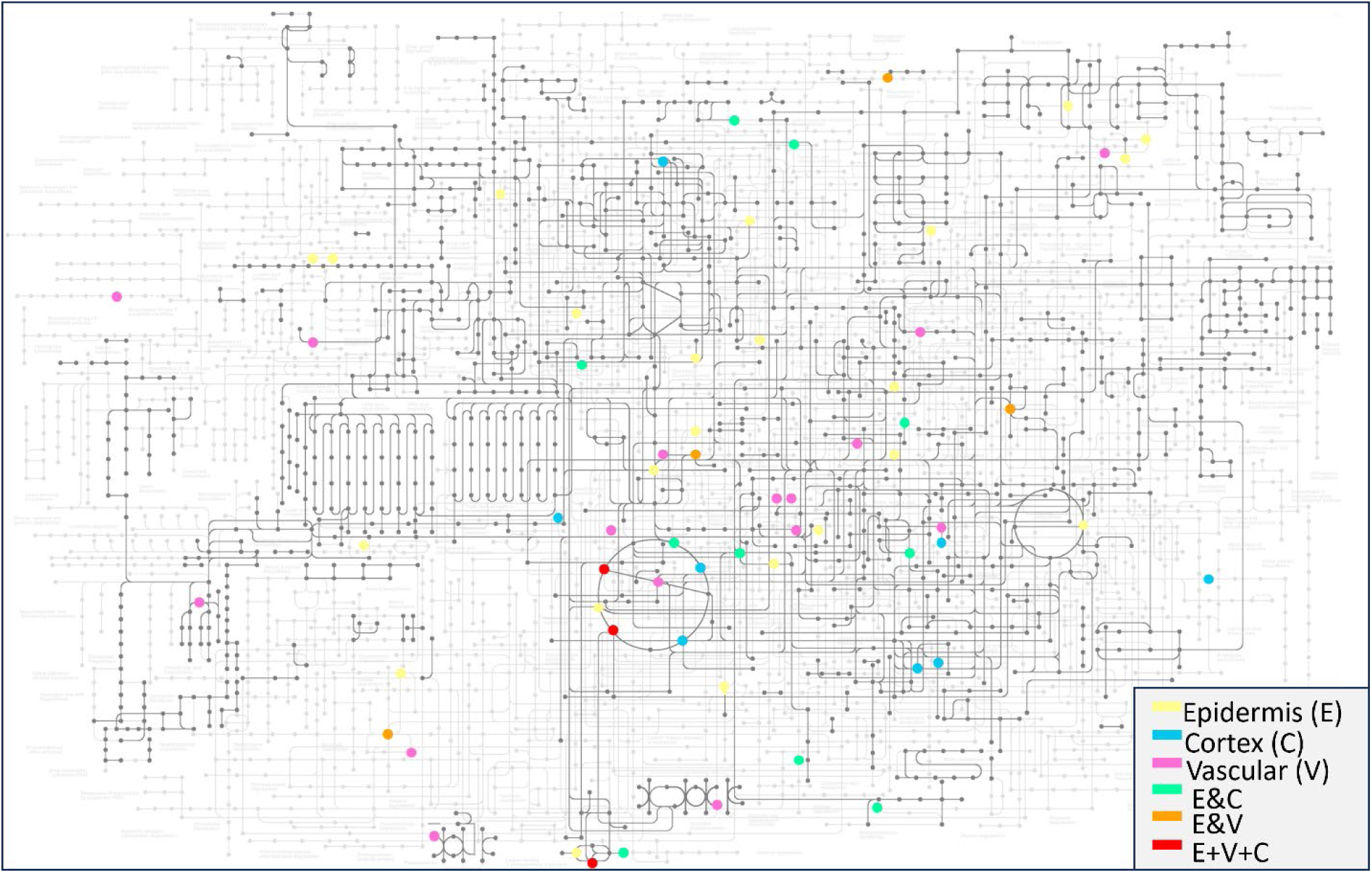
The full metabolic map for *Populus trichocarpa* shows which metabolites are differentially abundant between all inoculated plants (WWE and DE grouped together) and all control plants (DC and WWC grouped together). Colors indicate which cell types were significantly different in that metabolite according to the legend.

**Table S1:** Metabolic pathways which contain at least one metabolite with significantly higher abundance in droughted plants with endophytes compared to non-inoculated droughted plants (DC vs. DE treatments).

| Pathway Kegg ID | Pathway name | Metabolites Identified |
| --- | --- | --- |
| M00030 | Lysine biosynthesis | 2-Oxoglutarate<br>(R)-2-Hydroxybutane-1,2,4-tricarboxylate |
| map00190 | Oxidative phosphorylation | Succinate<br>Fumarate |
| M00373 | Ethylmalonyl pathway | Glyoxylate<br>Malate |
| M00433 | Lysine biosynthesis | 2-Oxoglutarate |
|  |  | • (R)-2-Hydroxybutane-1,2,4-tricarboxylate |
| M00539 | Cumate degradation | • Isobutyric acid<br>• 2-Hydroxy-2,4-pentadienoate |
| M00545 | Trans-cinnamate degradation | • 2-Hydroxy-2,4-pentadienoate<br>• Acetaldehyde |
| M00608 | 2-oxocarboxylic acid chain extension | • (R)-2-Hydroxybutane-1,2,4-tricarboxylate<br>• 2-Oxoglutarate |
| M00948 | Hydroxyproline degradation | • 2-Oxoglutarate<br>• 2,5-Dioxopentanoate |
| M00956<br>M00957<br>M00960 | Lysine degradation | • 2-Oxoglutarate<br>• Succinate |
| map00073 | Cutin, suberin and wax biosynthesis | • 16-Hydroxypalmitate (up)<br>• 16-Oxopalmitate (down) |
| map00270 | Cysteine and methionine metabolism | • 5'-S-Methyl-5'-thioinosine<br>• Methanethiol |
| M00012 | Glyoxylate cycle | • Glyoxylate<br>• Malate |
| M00022 | Shikimate pathway | • D-Erythrose<br>• Shikimate<br>• 3-Dehydroshikimate |
| M00346 | Formaldehyde assimilation, serine pathway | • Glyoxylate<br>• Malate<br>• Hydroxypyruvate |
| M00374 | Dicarboxylate-hydroxybutyrate cycle | • Succinate<br>• Fumarate<br>• Malate |
| M00569 | Catechol meta-cleavage | • 2-Hydroxy-2,4-pentadienoate<br>• Acetaldehyde<br>• Catechol |
| M00740 | Methylaspartate cycle | • Malate<br>• 2-Oxoglutarate<br>• Glyoxylate |
| M00982 | Methylcitrate cycle | • Fumarate<br>• Malate<br>• Succinate |
| M00011 | Citrate cycle, second carbon oxidation | • 2-Oxoglutarate<br>• Succinate<br>• Fumarate<br>• Malate |
| M00173 | Reductive citrate cycle (Arnon-Buchanan cycle) | • 2-Oxoglutarate<br>• Succinate<br>• Fumarate<br>• Malate |
| M00376 | 3-hydroxypropionate bi-cycle | • Malate<br>• Succinate<br>• Glyoxylate<br>• Fumarate |
| M00532 | photorespiration | • Glycolate<br>• Hydroxypyruvate<br>• 2-Oxoglutarate<br>• Glyoxylate |
| M00620 | Incomplete reductive citric acid cycle | • Fumarate<br>• Malate<br>• 2-Oxoglutarate<br>• Succinate |
| M00009 | Citric acid cycle | • Fumarate<br>• Malate<br>• 2-Oxoglutarate<br>• Succinate |

**Table S2:** *Features identified from Lasso Regularization (alpha = 0.1) as correlated to the presence or absence of endophytes. Asterisk (*) indicates a putative metabolite annotation*.

| Metabolite | Metabolite Location | Lasso Regularization Coefficient |
| --- | --- | --- |
| Succinate, Methylmalonate, Methyl oxalate, (3R)-3-Hydroxy-4-oxobutanoate* | Internal (Top root) | -0.09178 |
| 10-Hydroxydecanoic acid | Root Exudate | -0.08541 |
| 1-epi-Valienol 7-phosphate * | Internal (Top root) | -0.02151 |
| Phosphoagmatine* | Internal (Top root) | -0.00098 |
| 3,4-Dihydroxybenzoic acid | Root Exudate | 0.012926 |
| n-Octyl sulfate | Root Exudate | 0.075603 |
| L-(-)-Phenylalanine | Root Exudate | 0.154643 |

**Table S3:** *Features identified from Lasso Regularization (alpha = 0.1) as correlated to the presence or absence of drought stress. Asterisk (*) indicates a putative metabolite annotation*.

| Metabolite | Metabolite Location | Lasso Regularization Coefficient |
| --- | --- | --- |
| Arginine | Exudate | -0.21161 |
| 4-Sulfobenzoate* | Internal (Top Root) | -0.01767 |
| 2-Hydroxypalmitic acid | Exudate | -0.01275 |
| 11-O-Demethylpradinone I* | Internal (Top Root) | -0.01254 |
| 4-O-.beta.-Galactopyranosyl-D-mannopyranose | Exudate | -0.00613 |
| <u>Deacetoxycephalosporin C</u>* | Internal (Mid Root) | 0.011354 |
| Baicalein | Exudate | 0.011803 |
| 2-Oxo-6-methylthiohexanoic acid* | Internal (Top Root) | 0.024216 |
| Glutaric acid | Exudate | 0.030211 |
| <u>Sulfo jasmonate</u> | Exudate | 0.091149 |

**Supplementary Figure 3.**
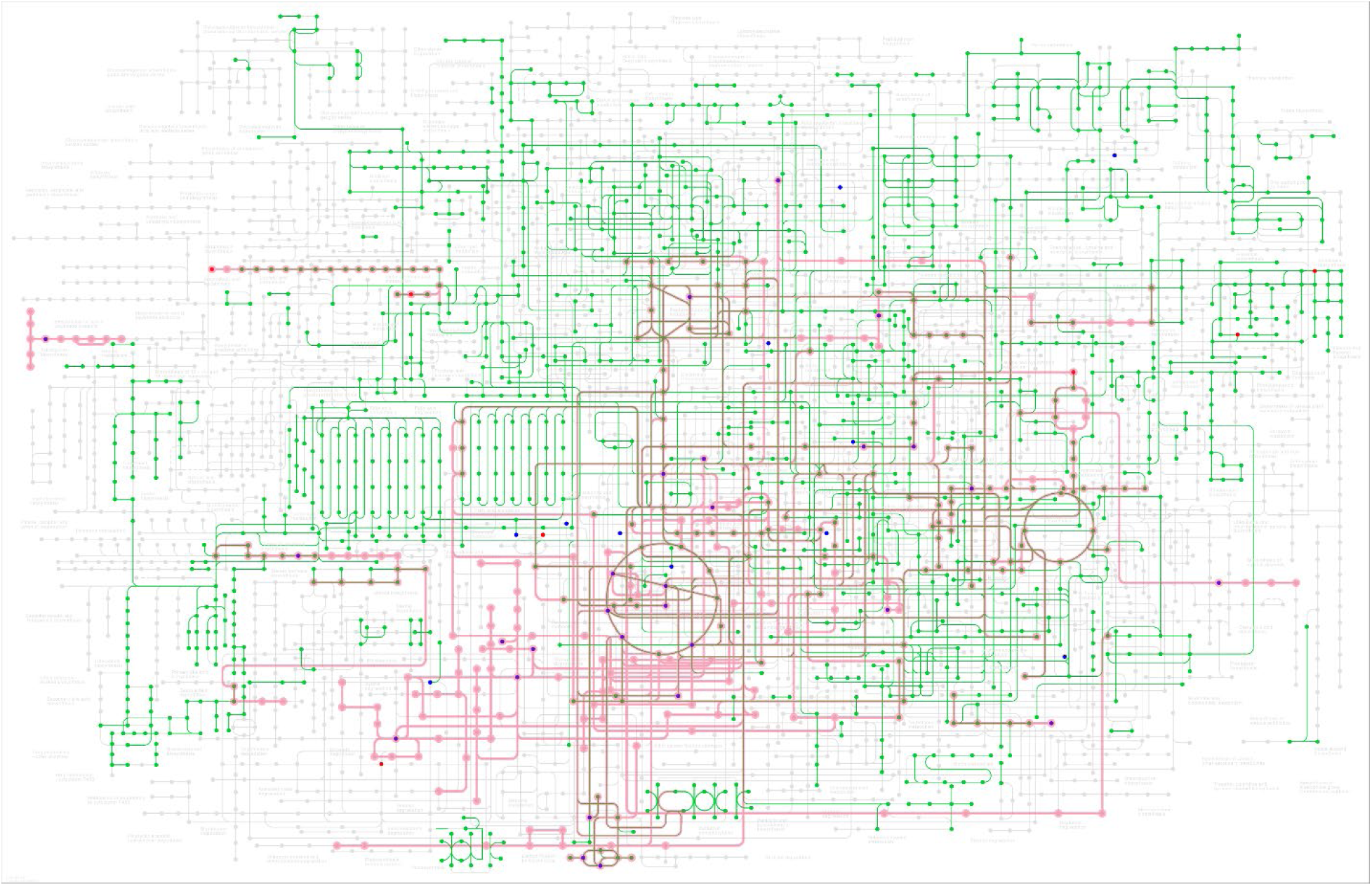
Map of all metabolites (blue or red dots) and metabolic pathways (pink) in all locations/cells and exudates which are significantly different between DC and DE conditions. Blue-colored dots are metabolites which were significantly more abundant when endophytes were added while red colored dots are metabolites that were significantly less abundant when endophytes were added. 89 potential metabolic pathways are affected by endophytes under drought.

**Supplementary Figure 4:**
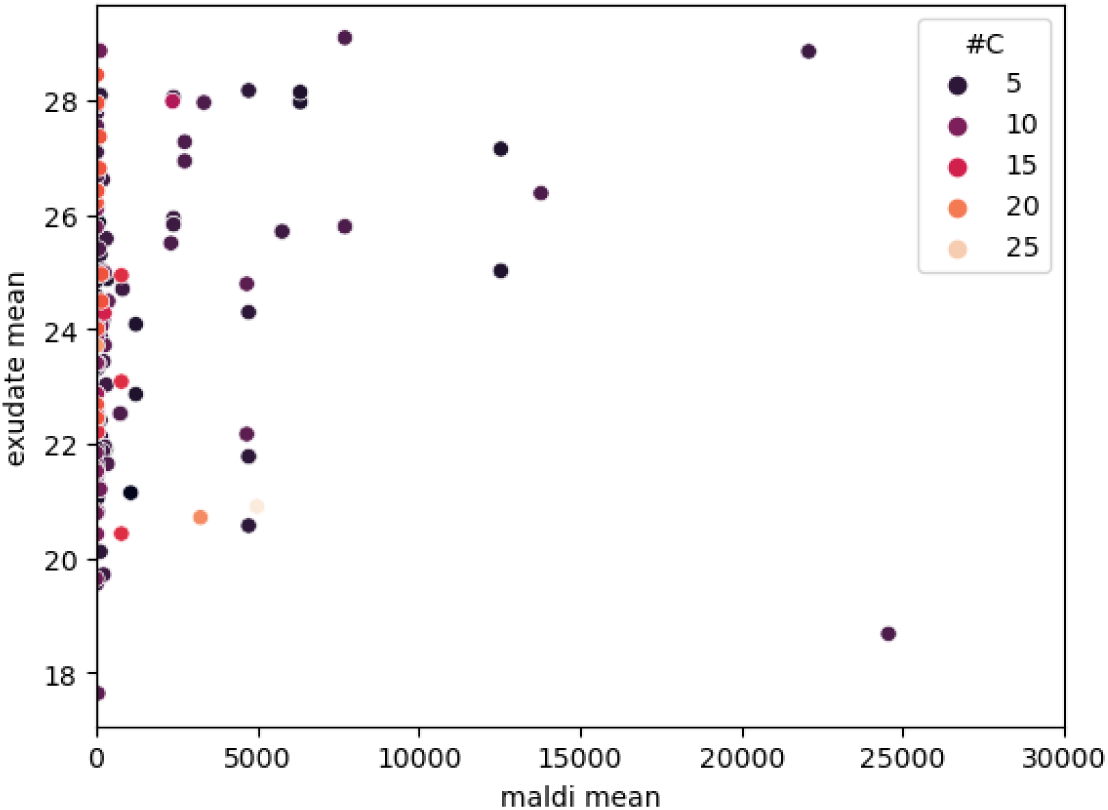
No correlation existed between the average abundance of metabolites within all root tissues and the average abundance of metabolite exuded by the root. In the above graph, each dot represents a metabolite and its mean value within the root is plotted on the x axis while its mean value in the exudate is plotted on the y value. Color indicates the number of carbon atoms that each metabolite contains.

**Supplementary Figure 5:**
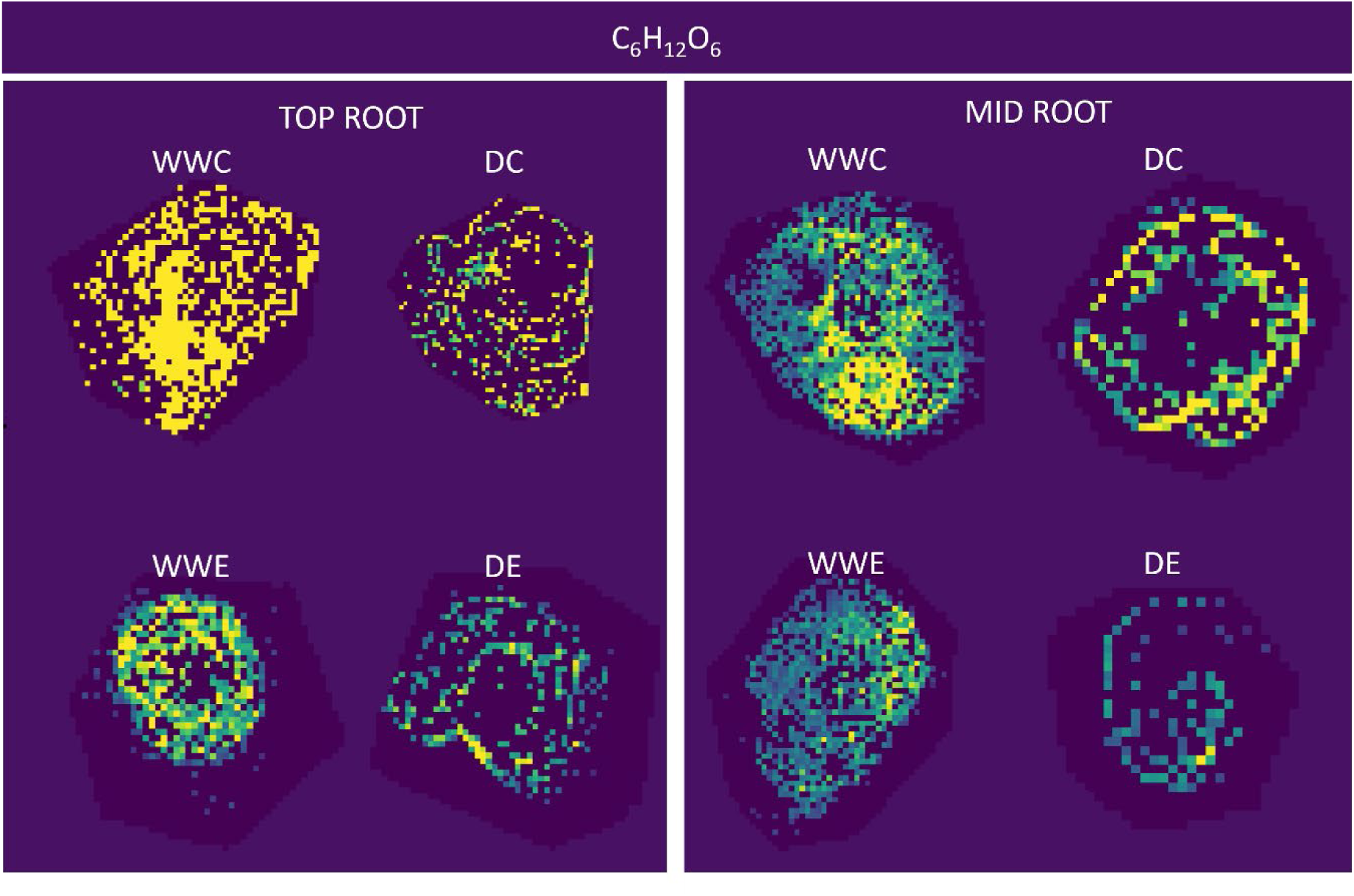
The relative abundance of C_6_H_12_O_6_ sugars (including glucose) is significantly reduced in droughted tissue compared to well-watered root tissue. Note that the MALDI MSI technique used here cannot discern between C_6_H_12_O_6_ sugar isomers. Representatives of root cross sections of top and mid root zones are shown. WWC: Well-watered control; WWE: Well-watered and endophyte inoculated; DC: Droughted control (DC); Droughted and endophyte inoculated (DE).

